# AI assisted Design of Ligands for Lipocalin-2

**DOI:** 10.1101/2025.05.18.654718

**Authors:** Jacopo Sgrignani, Sara Buscarini, Patrizia Locatelli, Concetta Guerra, Alberto Furlan, Yingyi Chen, Giada Zoppi, Andrea Cavalli

**Author notes:** Corresponding authors: Dr. Andrea Cavalli, Institute for research in Biomedicine (IRB), Universita’ della Svizzera Italiana (USI), Via Chiesa 5, 6500 Bellinzona (CH), Dr. Jacopo Sgrignani, Institute for research in Biomedicine (IRB), Universita’ della Svizzera Italiana (USI), Via Chiesa 5, 6500 Bellinzona (CH). These authors share first authorship (Jacopo Sgrignani, Sara Buscarini).

## Abstract

Lipocalin-2 (LCN2) is an acute-phase glycoprotein whose upregulation is linked to blood– brain-barrier breakdown and neuroinflammation, making it an attractive diagnostic and therapeutic target. We developed an end-to-end, AI-guided workflow to rapidly design *de-novo* miniproteins that bind LCN2. Backbone scaffolds were generated with RFdiffusion, sequences were optimized with ProteinMPNN, and candidates filtered *in silico* using a consensus of AlphaFold2 confidence metrics (mean interface pAE < 10) and binding free energy predicted by Prodigy. From an initial library of 10,000 designs, five were expressed and purified from *E. coli*. Using biolayer interferometry (BLI) we identified MiniP-2 as the lead construct, exhibiting a dissociation constant (Kd) of 4.2 nM. Structural modeling revealed that binding is primarily mediated by backbone hydrogen bonds along with a stabilizing salt bridge between Arg37 of MinP-2 and Asp97 of LCN2. These findings demonstrate that a fully computational generative workflow can yield nanomolar LCN2 binders in a single design–build–test cycle. MinP-2 represents a promising starting point for affinity maturation, structural studies, and *in vivo* evaluation as an imaging probe or antagonist of LCN2-mediated signaling. Specifically, SPR competition experiments showed that MinP-2 can inhibit LCN2 binding to MMP-9, suggesting its potential to mitigate the pathological effects of this interaction within the central nervous system.

## Introduction

Lipocalin-2 (LCN2) - also known as neutrophil gelatinase-associated lipocalin, siderocalin or 24p3 - is an acute-phase glycoprotein rapidly upregulated during infection and sterile inflammation (Chandrasekaran et al., 2024). Like other members of the lipocalin family, LCN2 adopts the canonical eight-stranded β-barrel, which allows it to bind bacterial siderophores and sequester iron, thereby acting as a key first-line defense mechanism against pathogens (Figure 1) (Chia et al., 2011;Chandrasekaran et al., 2024).

**Figure 1.**
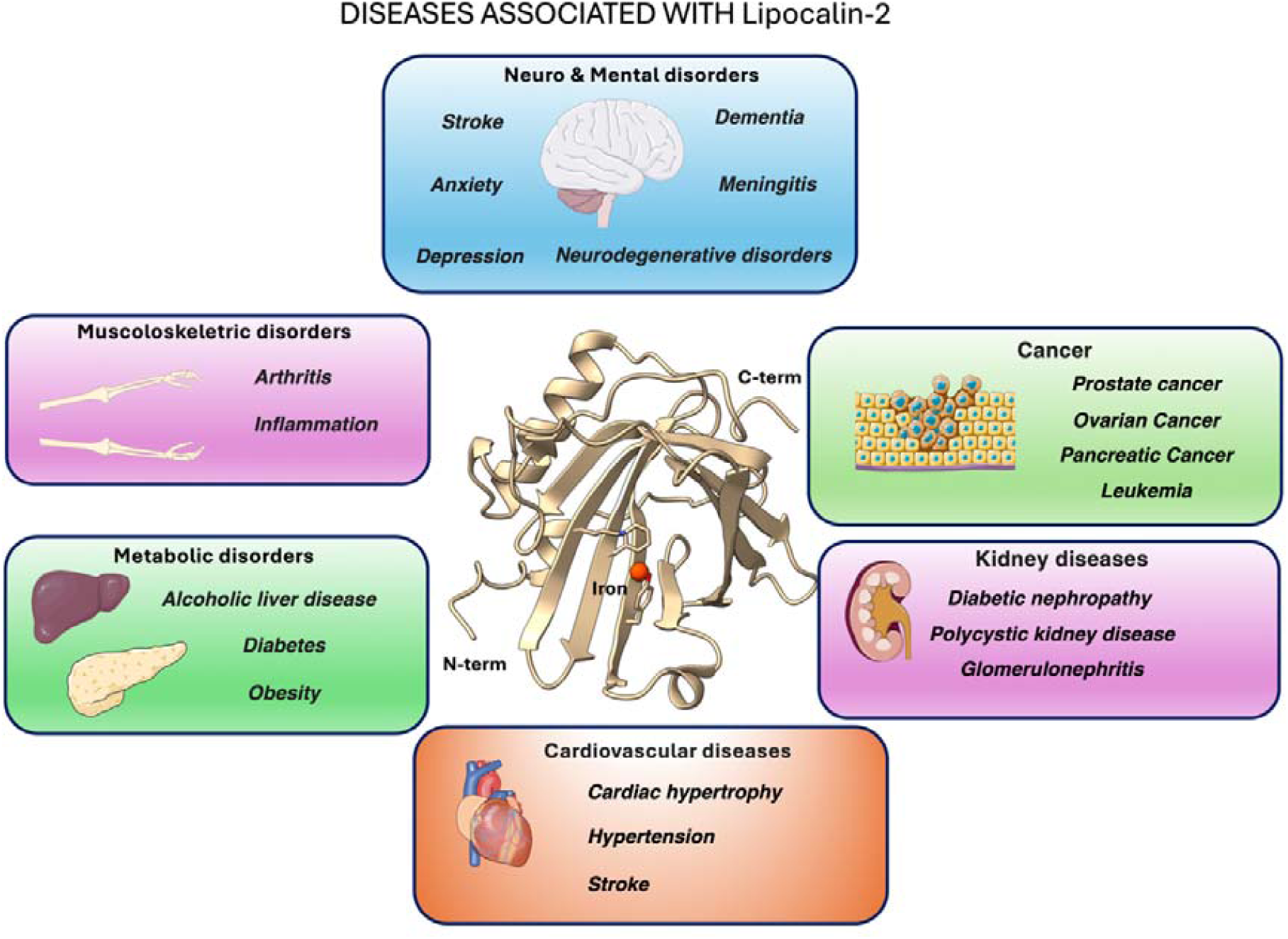
Graphical overview of major pathological conditions linked to LCN2 (lipocalin-2, also known as neutrophil gelatinase-associated lipocalin, siderocalin or 24p3) overexpression or activity. At the center, the X-ray crystal structure of human LCN2 (PDB ID: 3FW4; Bao et al., 2010). The images of the organs represented in the pictures were adapted from Servier Medical Art (https://smart.servier.com/), licensed under CC BY 4.0 (https://creativecommons.org/licenses/by/4.0/).

The interaction between LCN2 and its receptor NGALR plays a significant role in various cellular processes and is implicated in several pathological conditions. In particular, binding to NGALR amplifies pro-inflammatory cytokine release (Jha et al., 2015;Shao et al., 2016) and, when over-expressed by reactive glia, contributes to neurotoxicity in several central nervous system (CNS) disorders (Afridi et al., 2024).

LCN2 also forms stable heterodimers with MMP-9, a zinc-dependent matrix metalloproteinase (Triebel et al., 1992; Kjeldsen et al., 1993). This interaction stabilizes MMP-9, preventing its degradation. As a key enzyme for extracellular matrix remodeling, MMP-9 plays a critical role in breaking down extracellular matrix components. Excessive MMP-9 activity disrupts the integrity of the Blood-Brain Barrier (BBB), contributing to the development of neurodegenerative diseases such as Alzheimer’s and Parkinson’s (Jung and Ryu, 2023). Dysregulation of MMP-9 has also been linked to ischemia, trauma, neurodegenerative disorders (Ram et al., 2006) (Shigemori et al., 2006) and brain tumors (Vafadari et al., 2016).

Despite LCN2’s central role in neuroinflammation, no blood–brain barrier (BBB) permeable ligands exist that can either block the LCN2–NGALR signaling or disrupt the LCN2–MMP-9 heterodimer - both promising therapeutic strategies. Current therapeutic approaches against LCN2-driven pathogenic mechanisms aim to reduce its expression or secretion, enhance its degradation, block its activity with neutralizing antibodies, inhibit receptor binding, or interfere with downstream signaling pathways. Among small-molecule approaches, promising outcomes have been reported for the proteasome inhibitor bortezomib (Bae et al., 2024), which reduces LCN2 transcription, and for the autophagy activator Torin 1, which decreases LCN2 protein levels (Qiu et al., 2024). Antibody-based therapies represent another viable strategy to block LCN2 function and signaling. However, their large molecular weight severely limits blood–brain barrier (BBB) permeability, reducing their effectiveness in neurological applications. To overcome these limitations, we propose that engineered miniprotein binders (typically proteins with a mass below 10 kDa) with high affinity for LCN2 could serve as a versatile and BBB-compatible therapeutic platform. Protein-protein interaction (PPI) modulators are generally classified as small molecules, peptides, and antibodies (Nada et al., 2024). While small molecules offer advantages such as oral bioavailability and stability, peptides or antibodies are often more effective at disrupting PPIs, especially those involving large, flat interaction interfaces lacking well-defined binding pockets.

Recent advances in AI-driven protein design - pioneered by the Baker laboratory – have enabled the *de novo* generation of miniproteins (10–15 kDa) capable of binding targets of interest (Cao et al., 2020b; Huang et al., 2024; Weinberg et al., 2024;Yang et al., 2025).

Building on these advances, we established an AI-guided pipeline to design de novo miniproteins that block the LCN2 interfaces involved in NGALR and MMP-9 binding. Using this approach, we computationally designed, expressed in E. coli, and biophysically characterized five AI-generated candidates, identifying a lead binder with single-digit nanomolar affinity for LCN2.

## Methods

### General reagents and buffers

All chemicals were analytical grade. Recombinant human LCN2 was produced and purified by GenScript. The MMP-9 protein used in SPR studies was acquired from antibodies.com (code A331071-50). Twin-Strep–tag magnetic beads, Buffer W and Buffer BXT were from IBA LifeSciences. Dynamic-light-scattering (DLS) and nano-DSF assays were carried out in phosphate-buffered saline (PBS; 20 mM sodium-phosphate, 150 mM NaCl, pH 7.4). All other solutions are described below.

### Miniprotein computational design

The generation of miniproteins to bind LCN2 was carried out using the procedure described by Baker and co-workers (Watson et al., 2023) (https://github. com/RosettaCommons/RFdiffusion). The crystal structure of LCN2 (PDB ID: 1DFV) was selected as template structure (Goetz et al., 2000). Following the protocol author’s suggestion, we generated 10,000 backbone scaffolds using RF diffusion (Watson et al., 2023) and subsequently designed 10,000 sequences with ProteinMPNN (Dauparas et al., 2022) to create LCN2 binders. Finally, the ability of the generated sequences to correctly fold and to bind the target was evaluated using AlphaFold2 (AF2) (Jumper et al., 2021).

For each model we recorded the mean pLDDT and the interface predicted aligned-error (pAE_interface). Designs with pAE_interface<10 were retained, yielding 1,184 candidates. Next, to select the five most promising binders, we estimated the LCN2-binder affinity with PRODIGY, a contact-based predictor of protein-protein affinity (Vangone and Bonvin, 2017). Finally, the five proteins reporting the most negative ΔG were expressed and purified.

### Alanine scanning with DrugScorePPI server

To identify the molecular determinants of the MinP-2/LCN2 interaction, we analyzed the AF2-predicted complex structure using the DrugScorePPI server (Kruger and Gohlke, 2010).

### Assessment of protein expression

The DNA sequences encoding the five select miniproteins were synthesized and cloned by GenScript into the pET30b plasmid, incorporating a Twin-Strep tag for purification. *E. coli* BL21 (DE3) pLysS cells were transformed with the plasmids and plated onto five separate LB-agar plates. Following overnight incubation, colonies that successfully incorporated the plasmids were selected.

Before starting the protein production and purification, we assessed miniprotein expression in *E.coli* BL21 (DE3) pLysS cells analyzing induced and not induced cultures by Western-Blot (WB).

Samples (20 µl) from both IPTG-induced and non-induced cultures were mixed with 20 µl of sample buffer containing Tris-Glycine SDS and a sample reducing agent. From these prepared stocks, 20 µl were loaded onto the gel. Following electrophoresis, the SDS-PAGE gel was transferred into a membrane. After the transfer, the membrane was activated with methanol and subsequently washed with Tris-buffered saline containing 0. 1% Tween® 20 (TBST). Following the washing step, blocking was performed with TBST and milk to prevent nonspecific binding. Subsequently, another washing step was conducted, and the primary antibody (IBA Lifesciences 2-1507-001; Anti-Strep) was added.

The next day, the primary antibody was removed, and the secondary antibody (anti-mouse IgG1 conjugated with horseradish peroxidase (HRP)) was applied. After the incubation step, another washings step was performed, and reagents were added to develop the Western Blot.

### Miniprotein expression and purification

Following positive WB reults, a bacterial preculture was prepared by inoculating a single colony into 7 mL of Luria-Bertani (LB) medium supplemented with chloramphenicol and kanamycin, and incubated at 37°C to promote bacterial growth.

Subsequently, 1 mL of the preculture was inoculated into 200 mL of LB medium and incubated at 37□°C until the culture reached an OD□□□ of 0.6. Protein expression was then induced by adding IPTG to a final concentration of 1 mM. After overnight incubation, bacterial cells were resuspended in a PBS added with two tablets of protease inhibitor (cOmplete™, Mini, EDTA-free Protease Inhibitor Cocktail) and lysed by sonication. The lysate was then filtered through a 0.22 μm membrane to remove debris before purification.

Strep-tag purification was performed using magnetic beads (IBA LifeSciences). Beads were washed with 1x Buffer W, followed by an incubation step to allow protein binding. After additional washing, elution was carried out using 1X Buffer BXT, yielding purified miniproteins in the eluate. Residual biotin in the elution buffer was removed by dialysis. Purity was verified by SDS-PAGE.

### Dynamic-light scattering (DLS) and nano-DSF

The size distribution and thermal stability of all miniproteins were analyzed using a Prometheus Panta (NanoTemper). Measurements were performed in triplicate, with miniproteins directly loaded into capillaries at concentrations of 0.32, 0.22, 0.18, 0.10, and 0.2 mg/mL for MinP-1 to MinP-5, respectively.

### Bio-layer interferometry (BLI)

Interactions between the designed miniproteins and LCN2 were analyzed using biolayer interferometry (BLI). LCN2 was immobilized by His-tag on BLI sensors Octet® Ni-NTA (NTA) Biosensors using a concentration of 5μg/ml. All the experiments were run using a Octet® R8 instrument. All miniproteins were tested at a concentration of 4 μM. For MinP-2, which showed clear binding, titration experiments were performed at 1000, 500, 250, 125, 62.5, 31.2, and 15.6□nM. The signal from a reference sensor without immobilized protein was subtracted to remove background noise and aspecific binding. Association and dissociation phases were 300 s each, and data were fitted globally with a 1:1 and 2:1 binding models in the Octet Analysis software.

#### Surface plasmon resonance (SPR)

SPR experiments were performed using a Biacore 8K system (Cytiva, Marlborough, MA, USA). MinP-2 dissolved in 10 mM sodium acetate (pH 4.5) at a concentration of 2 μM was immobilized via amine coupling onto a CM5 sensor chip (GE Healthcare, Chicago, IL, USA). MMP-9 was immobilized using the same procedure using protein concentration of 300 nM.

Five concentrations of LCN2 (7.8, 15.6, 31.2, 62.5, and 125 nM) were injected over the immobilized MinP-2 at a flow rate of 30 μL/min. The contact time for each injection was 120 s, while the dissociation phase was run for 600 s. All the experiments were performed at 25 °C. Following blank subtraction, the binding data were fitted and analyzed using Biacore Insights Evaluation Software v. 5.0.18. In the case of the MMP-9/LCN2 interaction a single concentration of 200 nM LCN2 alone or mixed with an equimolar concentration of MinP-2 was injected for 45s followed by 60s of dissociation.

## Results

### In-silico miniprotein design pipeline and candidate selection

In the initial phase of the project, we applied the computational pipeline originally proposed by Baker and colleagues (Watson et al., 2023). Starting from 10,000 backbone scaffolds generated with RFdiffusion, a single ProteinMPNN sequence was designed for each scaffold. AlphaFold2 rescoring retained 1,184 binders with a pAE_interaction score below 10 - a threshold shown to enrich for functional hits (Bennett et al., 2023) (Figure 2A)1S.

**Figure 2.**
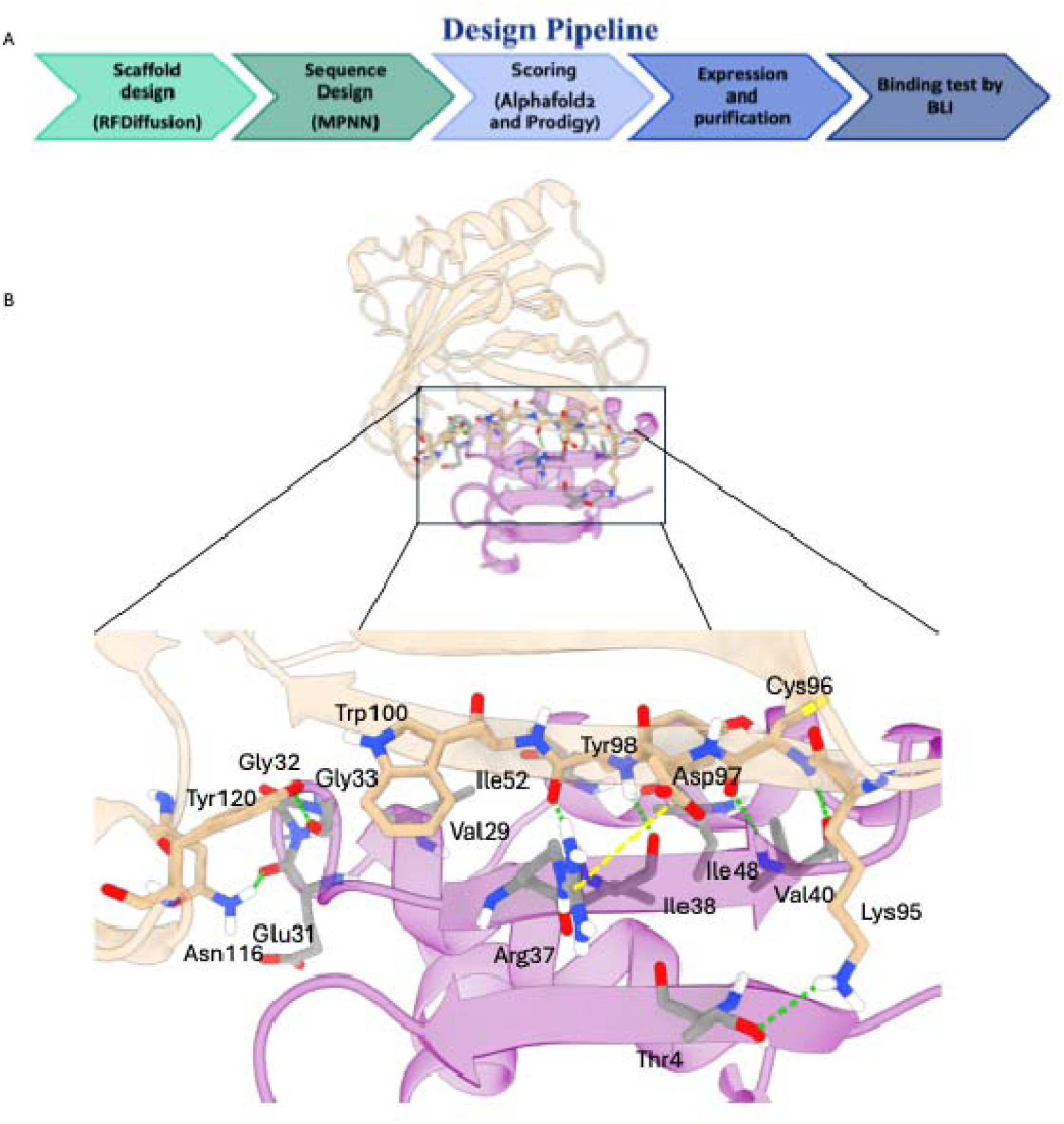
(A) Schematic representation of the applied computational pipeline. (B) Key molecular interactions driving the binding of MinP-2 (magenta cartoons) to LCN2 (white cartoons). The key interactions highlighted by the black box are shown in detail in the lower panel. Hydrogen bonds are shown as green dotted lines with the salt bridge as a dotted yellow line.

However, expressing and purifying such a large number of proteins is impractical. To overcome this limitation, we applied a consensus scoring approach to refine the selection. Specifically, all 1,184 candidates were re-ranked with the contact-based affinity predictor PRODIGY (Vangone and Bonvin, 2017), and the five miniproteins with the most negative predicted ΔG were selected for expression (Table 1, Figure S3).

**Table 1.**
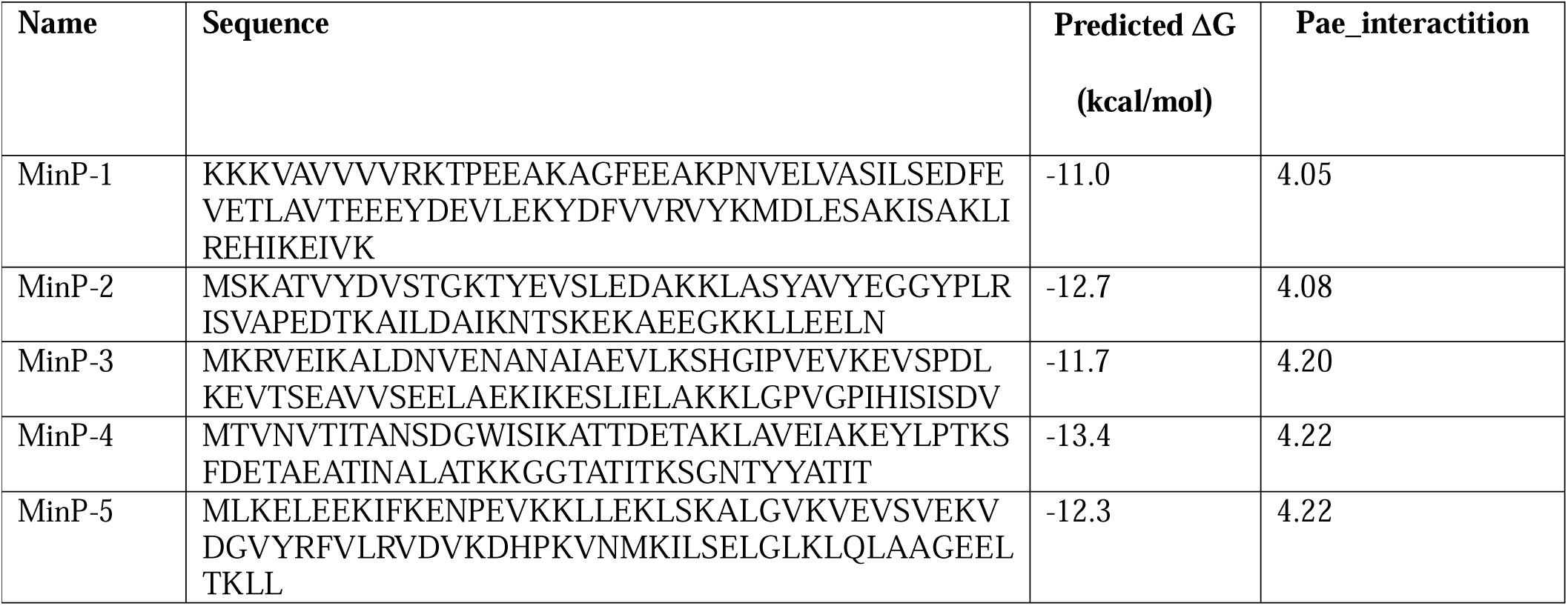
Aminoacidic sequences and predicted ΔG for the five miniproteins selected for further studies after computational design. The ligands are ordered by pae_interaction score.

### Expression, purification and biophysical characterization of AI designed miniproteins

Given that the proteins were designed by AI, before large-scale production we assessed the expression of the five constructs in *E. coli* by WB (Figure S1). This preliminary analysis confirmed that all proteins were expressed, although MinP-2 and MinP-3 showed lower expression levels. Consequently, we proceeded with large-scale expression and purification of all five constructs. Purification was performed using magnetic beads functionalized with a modified streptavidin (Strep-Tactin®XT, IBA Lifesciences), yielding highly pure proteins in sufficient quantities for subsequent biophysical characterization (Figure S2).

Before evaluating LCN2 binding, all purified constructs were characterized for size distribution via dynamic light scattering (DLS) and thermal stability via nanoDSF. DLS analysis revealed that all constructs - except MinP-4 - were monodisperse with an apparent hydrodynamic radius of approximately 2 nm, consistent with their predicted molecular weight and structural models (Figure 3-4). In contrast, MinP-4 showed a polydispersity index (PDI) of 0.38, indicative of aggregation. Thermal stability analysis by nanoDSF showed that only MinP-5 displayed a clear melting temperature (Tm at 46.8□°C). This is notably lower than the high thermal stability (Tm >□90□°C) reported for some AI-designed miniprotein binders by Baker and colleagues (Cao et al., 2020a).

**Figure 3.**
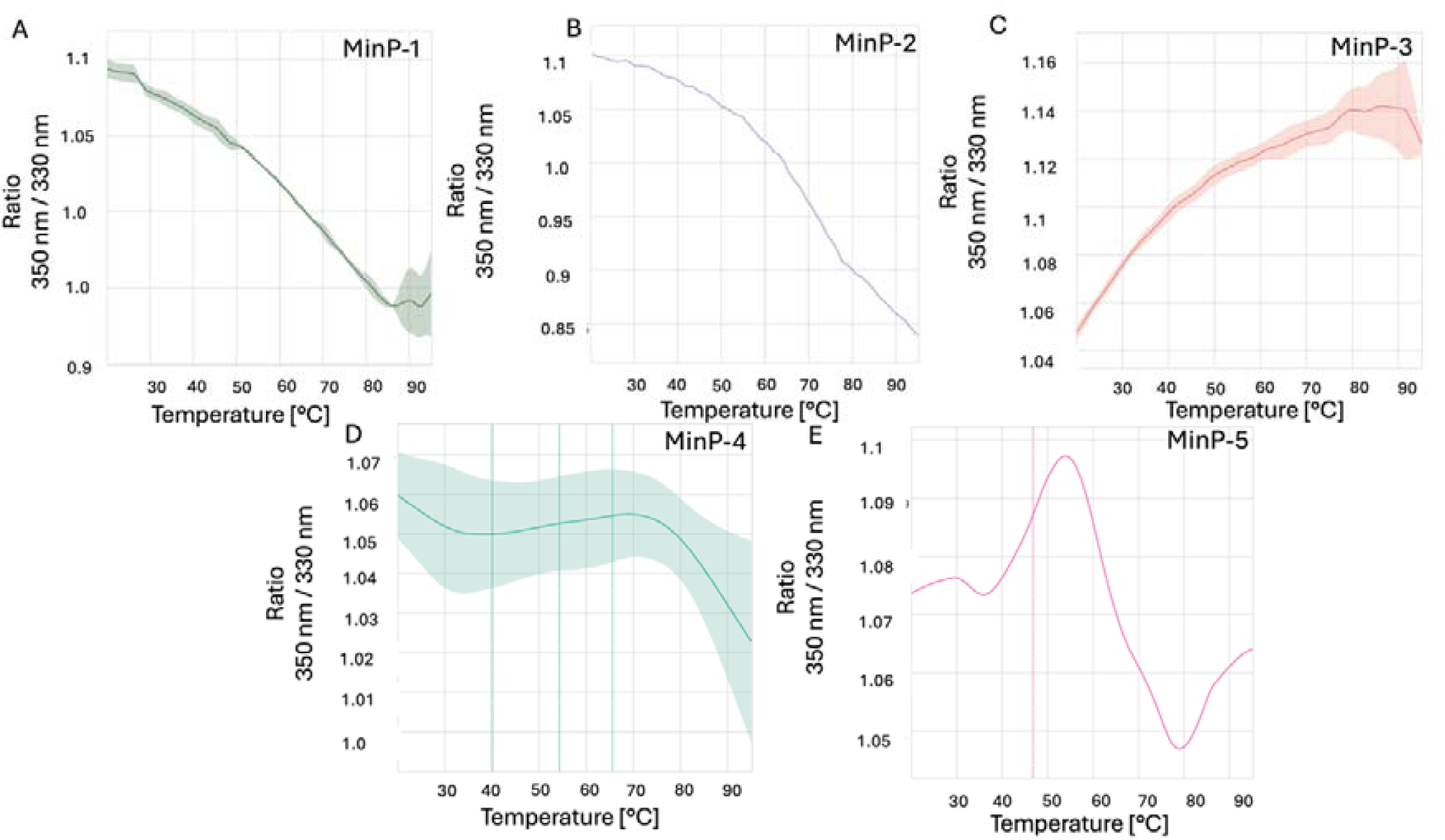
Results of nanoDSF experiments. The ratio of fluorescence intensity measured at 350 nm to the fluorescence intensity measured at 330 nm at different increasing temperatures for the five designed miniproteins This value reflects changes in the environment of tryptophan (Trp) residues within the protein making possible to monitor its stability at different temperatures.

**Figure 4.**
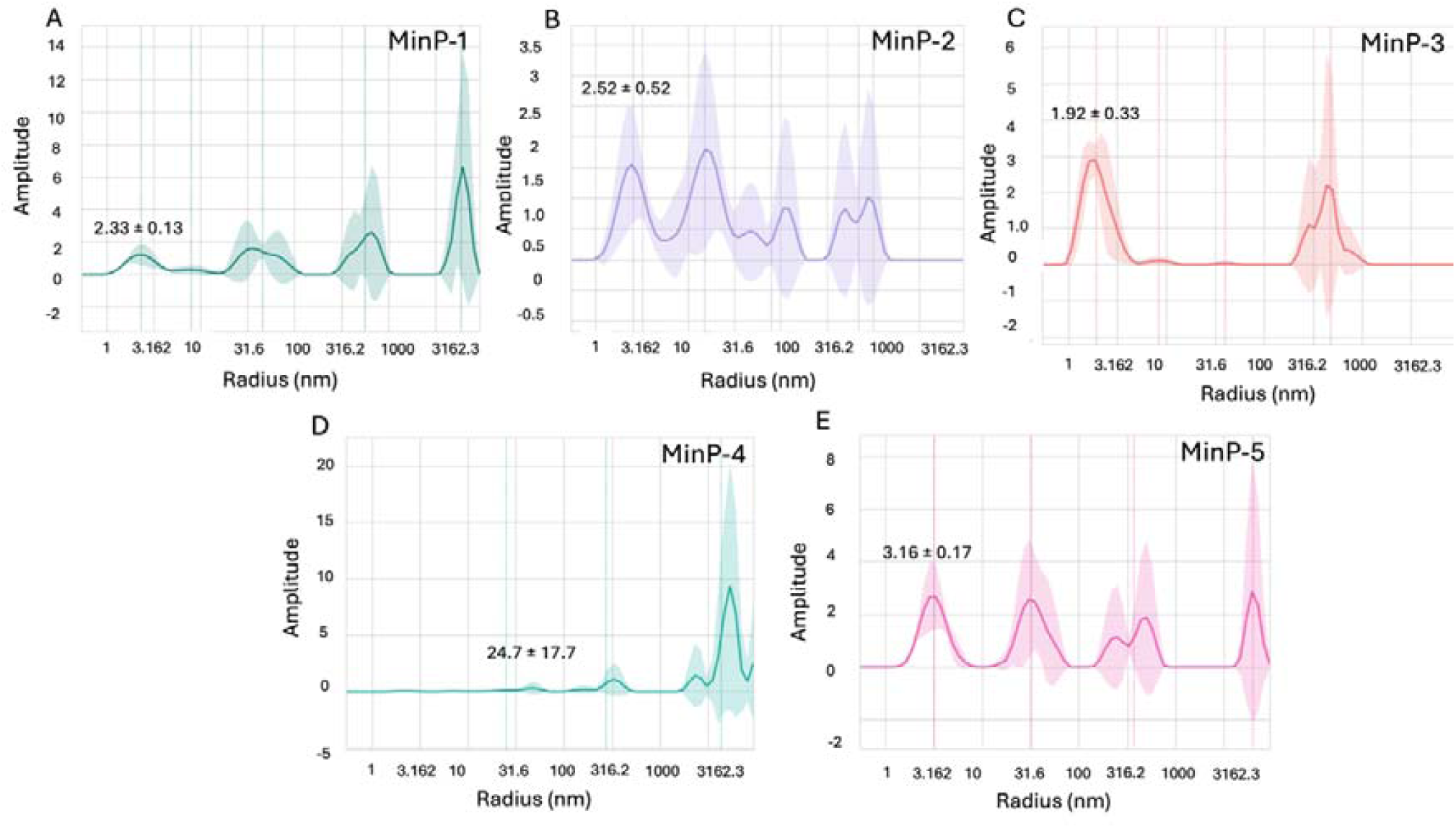
Results of DLS experiments. The DLS plots reflect the intensity of scattered light, which is proportional to the sixth power of particle size (∝ size□). Thus, larger particles appear overrepresented in intensity despite being fewer in number. For example, the ∼2□nm peak contains ∼2×10□ more particles than the ∼40□nm peak, and ∼2×10¹² more than the ∼400□nm peak, even though the peaks may appear similar in heigh.

**Figure 5.**
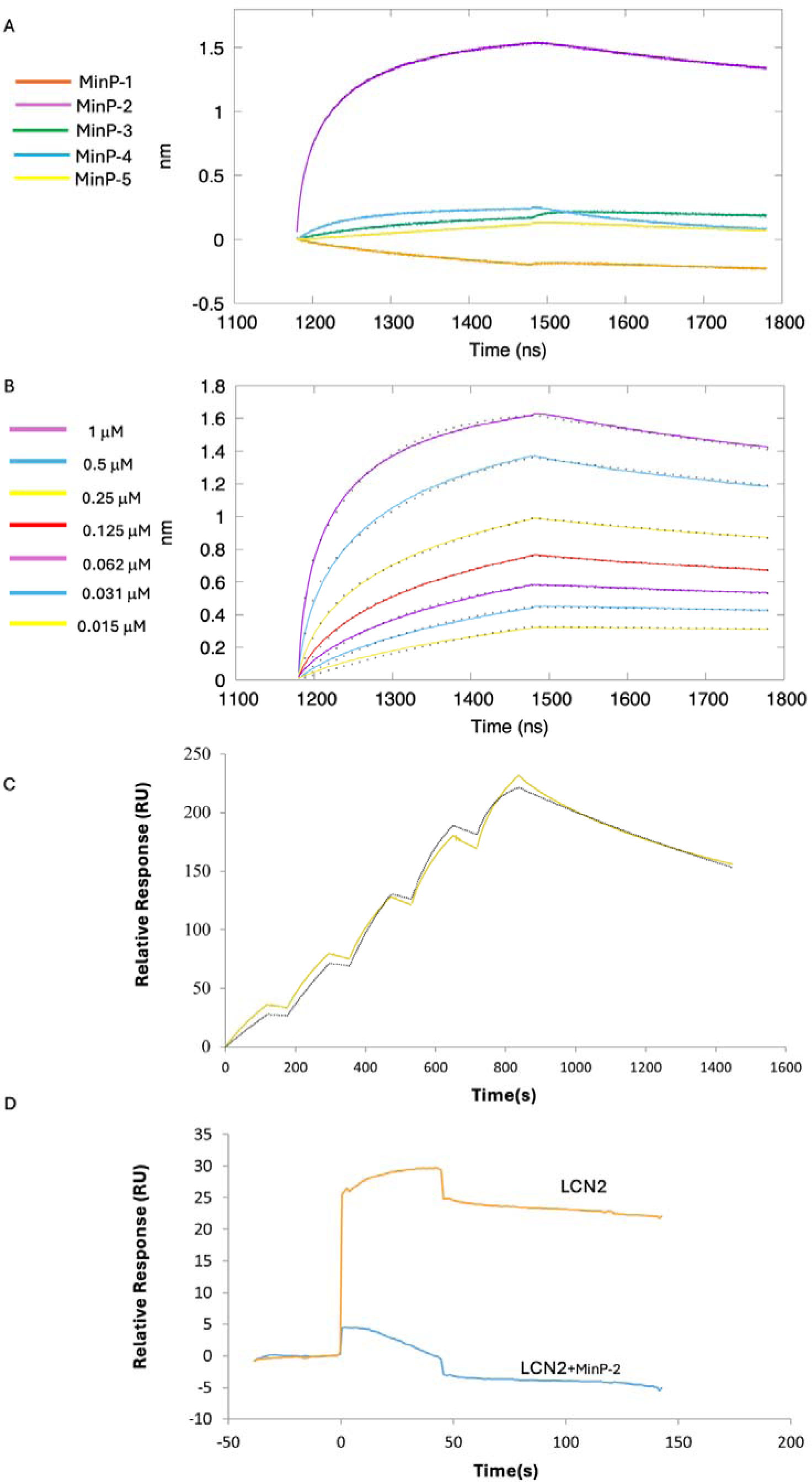
(A) Preliminary BLI analysis of miniprotein binding to LCN2 at a fixed concentration of 4 μM. Color coding: MinP-1 (orange), MinP-2 (violet), MinP-3 (green), MinP-4 (light blue), MinP-5 (yellow). (B) BLI titration curves of MinP-2 binding to LCN2 across seven concentrations, from top to bottom: 1 μM (violet), 0.5 μM (light blue), 0.25 μM (yellow), 0.125 μM (red), 0.0625 μM (violet), 0.0312 μM (light blue), and 0.0156 μM (yellow). Experimental data are shown as colored lines, with corresponding fitted values indicated by black dots. (C) Results of the LCN2/MinP-2 binding experiments by SPR. Sensogram is shown as a continuous yellow line, while fitted data are indicated as black dotted line. (D) Results of the SPR competition assay showing that MinP-2 inhibits MMP-9/LCN-2 binding.

**Figure 6.**
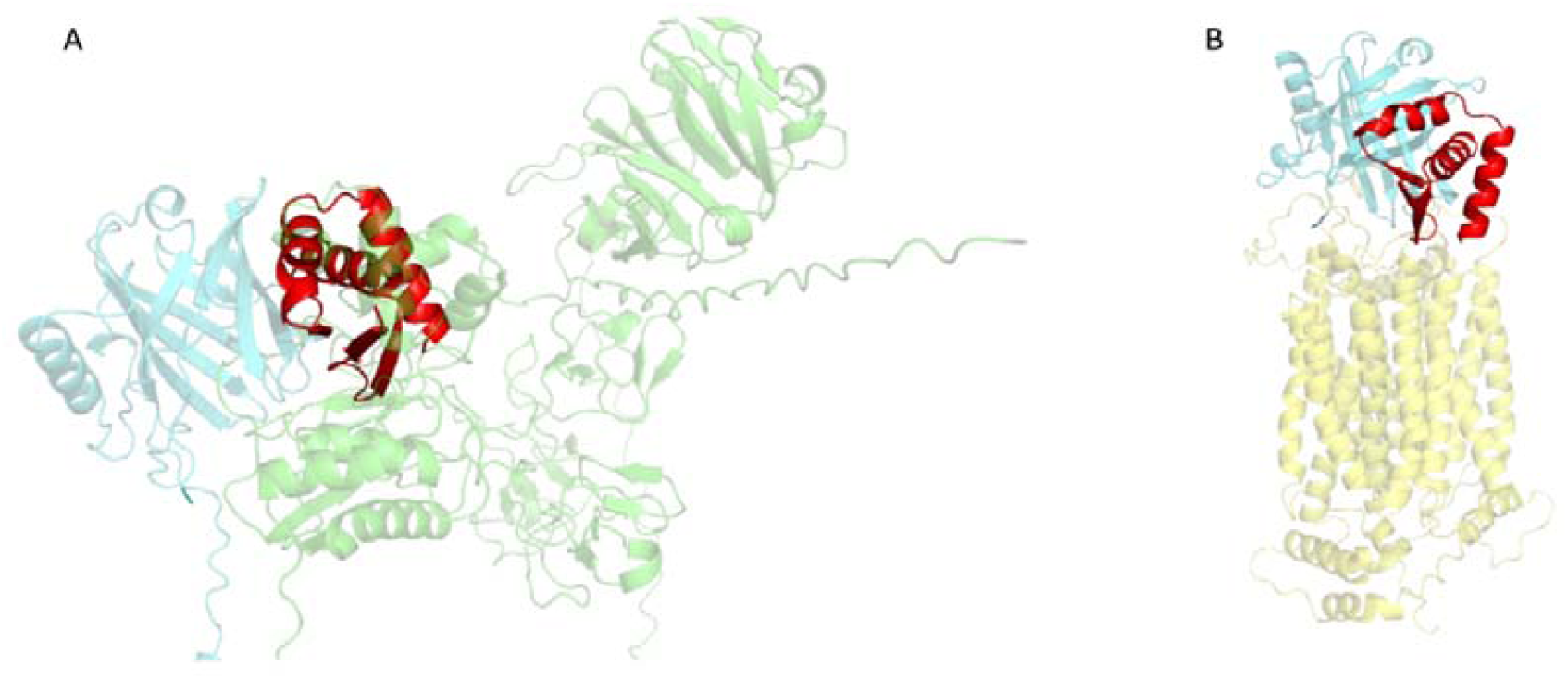
Complexes of LCN2 with MMP-9 and NGALR predicted by AlphaFold3, superimposed to LCN2/MinP-2. (A) Model of the LCN2 (aquamarine)-MMP-9 (green) complex, overlaid MinP-2 (red) structure. (B) Model of the LCN2 (aquamarine)-NGALR (yellow) complex, overlaid with the LCN2/MinP-2 (red). The images were produced by superimposing the LCN2 structure found in the MMP-9/LCN2 and NGALR/LCN2 complexes onto the LCN2 structure from the LCN2/MinP-2 complex. To enhance clarity, only the LCN2 structures from the MMP-9/LCN2 and NGALR/LCN2 complexes are displayed. These AlphaFold3-based superpositions indicate that MinP-2 can competitively bind LCN2 by occupying binding sites shared with both MMP-9 and NGALR.

### Binding of miniproteins to LCN2 investigated by BLI and SPR

Preliminary binding assays at 4□µM were performed to evaluate the interaction of the five miniproteins with LCN2. Among them, MinP-2 emerged as the most promising binder (Figure□3A), whereas the other constructs displayed only weak or negligible binding.

Therefore, to fully characterized affinity and kinetics of the LCN2-MinP-2 interaction we performed a titration across seven concentrations (Figure 3B). This experiment showed that MinP2 strongly binds LCN2, with two dissociation constants (Kd1 = 0.7 and Kd2 = 35 nM) obtained using a 2:1 heterogeneous model. The requirement for a 2:1 model may stem from surface crowding on the sensor, partially blocking MinP-2 binding sites and resulting in a secondary, lower-affinity interaction.

To verify if the immobilization strategy used in the BLI experiment could be the source of the 2:1 binding, we performed surface plasmon resonance (SPR) experiments (Figure 3C). In this case, confirming our hypothesis about the influence of the immobilization strategy on the results, the data were best fitted with a 1:1 model, yielding a Kd for the LCN2/MinP-2 complex of 4.2 nM.

### Structural insight into the LCN2/MinP-2 interface

To better understand the molecular determinants of the LCN2/MinP-2 interaction, we performed a computational alanine scanning analysis on the structure AlphaFold2-predicted complex using the DrugScorePPI server. The results (Tables S1-2) and visual inspection of the interface (Figure 2B) reveled that binding is determined by multiple intermolecular interactions, notably: (i) backbone hydrogen bonds between LCN2 residues Cys96, Asp97, and Tyr98 and the MinP-2 residues Ile38 and Val40 forming a short antiparallel β-augmentation interface; (ii) additional hydrogen bonds between LCN2 Lys95, Asn116, and Tyr120 with MinP-2 residues Thr4, Glu31, and Gly33; and (iii) a salt bridge between Arg37 of MinP-2 and Asp97 of LCN2. Taken together, these interactions rationalize the observed low-nanomolar affinity.

### Assessing the inhibition of the MMP-9/LCN2 interaction by SPR

To evaluate whether MinP-2 can prevent the interaction between MMP-9 and LCN2, we performed an SPR experiment in which LCN2 - alone or in presence of an equimolar (200 nM) concentration of MinP-2 - was injected over immobilized MMP-9. This experiment (Figure 3D) demonstrated that MinP-2 effectively LCN2/MMp-9 interaction.

## Discussion

Lipocalin-2 (LCN2) drives, among other effects, BBB breakdown and neuroinflammation through interactions with its partners NGALR and MMP-9. However, no ligands with a molecular weight smaller than an antibody are currently available to block these interactions. Here we show that a 12-kDa miniprotein (MinP-2), generated in a single AI-guided cycle, binds LCN2 with single-digit nanomolar affinity (Kd 4.2 nM). This affinity is comparable to that observed for monoclonal antibodies, but achieved with a 10-fold smaller molecule. Our strategy, incorporating a contact-based scoring approach, demonstrates that consensus rescoring with AlphaFold2 and PRODIGY facilitates the identification of true binders among a large number of AI generated miniproteins, thereby reducing experimental efforts in protein purification and binding tests. All five selected miniproteins were successfully expressed in *E. coli,* and four were monodisperse and thermostable.

In fact, quality control analyses performed using DLS and nanoDSF revealed that, with the exception of MinP-5, all proteins are thermally stable and predominantly monomeric in PBS at a concentration of ∼10–12 µM, consistent with previous reports (Vázquez Torres et al., 2024). However, MinP-4 exhibited a tendency toward aggregation, representing an exception to this general behavior.

From preliminary BLI experiments, MinP-2, a designed miniprotein, emerged as a promising candidate, demonstrating significant interaction with LCN2 and warranting further detailed characterization. Binding analysis revealed that MinP-2 interacts with LCN2 with a dissociation constant (Kd) in the low nanomolar range of 4.2 nM, underscoring its potential for future applications in therapeutic or diagnostic settings.

When comparing the success rate of our protocol to previous studies, the identification of a strong LCN2 binder among just 5 miniproteins is notable. For many target - such as TrkA, EGFR, InsulinR, CD3 (Cao et al., 2022) – hit rates are below 1% when a 400 nM affinity threshold, with 15,000 and 100,000 candidates tested. However, we acknowledge that the high success rate observed in our study might be influenced by the specific nature of LCN2 as a target. More extensive benchmarks are required to fully evaluate the impact of incorporating physical or contact-based scores into the binder selection process.

To better understand MinP-2’s binding mode, we examined AlphaFold2 models of the LCN2 MinP-2 complex. The analysis indicates that affinity is mediated by a short β-augmentation segment: backbone-to-backbone hydrogen bonds form between LCN2 and MinP-2 residues, creating a continuous strand that anchors the miniprotein to its target complemented by additional hydrogen bonds and a salt bridge between Arg37 of MinP-2 and Asp97 of LCN2.

To explore whether MinP-2 can block interactions with MMP-9 and NGALR, we predicted the structures of the respective complexes using AlphaFold3. The result of these calculations (Figure 4) suggested that MinP-2 has the potentiality to hinder their interaction with LCN2.

MMP-9 can be readily expressed, purified, and is commercially available. Therefore, to experimentally confirm our hypothesis, we carried out a competitive SPR experiments (Figure 4D). The results confirmed that MinP-2 can prevent the MMP-9/LCN2 interaction. Differently, NGALR is a membrane protein which expression and purification in a binding-competent form is challenging, therefore additional work will be necessary to experimentally prove

## Conclusions

In this work, we demonstrated that a fully automated, AI-driven workflow – combining RFdiffusion for backbone creation, ProteinMPNN for sequence optimisation, and AlphaFold2 for structure-based triage - can deliver potent miniproteins capable of binding LCN2 with high affinity.

From an initial library of 10,000 *in-silico* designs, five candidates were produced recombinantly, and one lead binder (MinP-2) displayed nanomolar affinity for LCN2 while maintaining high solubility and thermal stability. Furthermore, structural models suggest that binding is dominated by a short antiparallel β-augmentation interface, a motif that could be adapted or grafted onto other scaffolds to engineer new binders.

These findings establish a starting point for developing LCN2-targeted diagnostics or therapeutics that modulate neuroinflammatory pathways. Future work will focus on solving a high-resolution structure of the MinP-2–LCN2 complex and comprehensively evaluating the miniprotein’s ability to disrupt pathological LCN2 interactions in cellular and *in-vivo* models. More broadly, our study underscores the power of generative deep-learning methods to accelerate ligand discovery for challenging protein–protein interfaces, even in the absence of prior binder templates.

## Founding

This work was supported by the SNF grant CRSII—222762.

