## Supplementary material for "AI assisted Design of Ligands for Lipocalin-2"

**SUPPORTING MATERIAL**

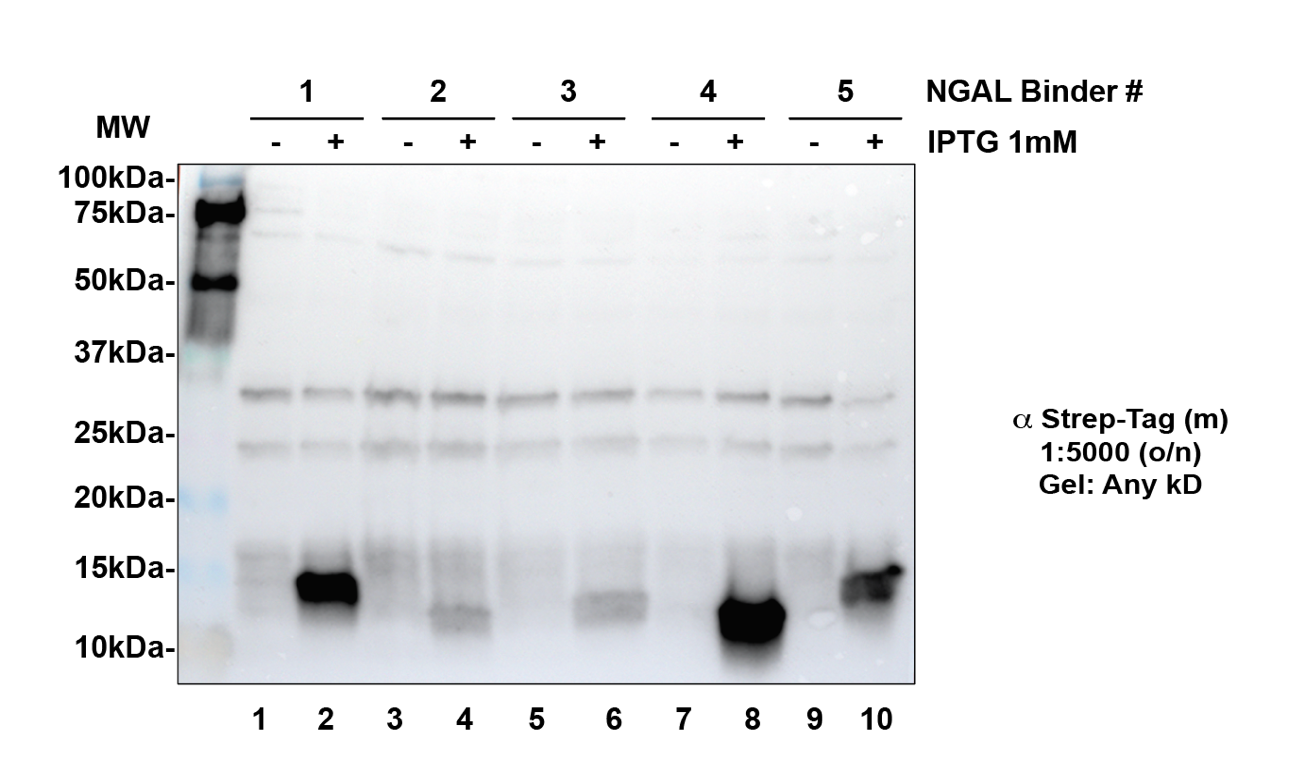

**Figure S1. Western Blot (WB) executed to assess the expression of the AI designed mini proteins prior large expression and purification**. For all the proteins bacterial cultures of induced and not induced bacteria were analyzed.

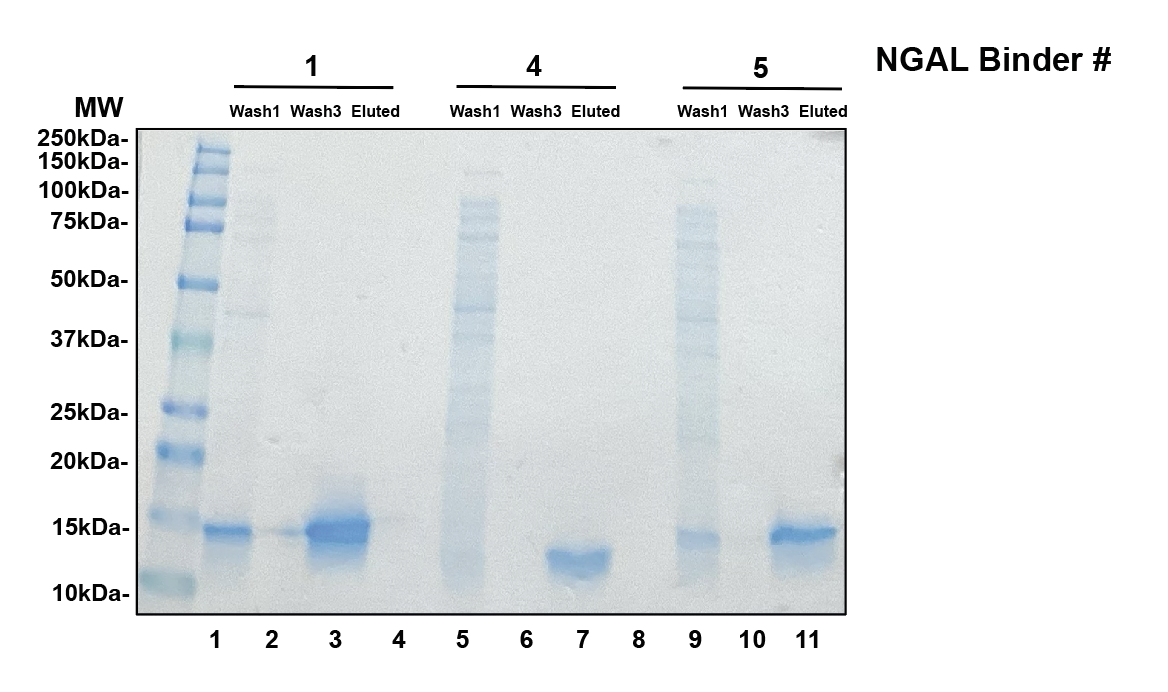

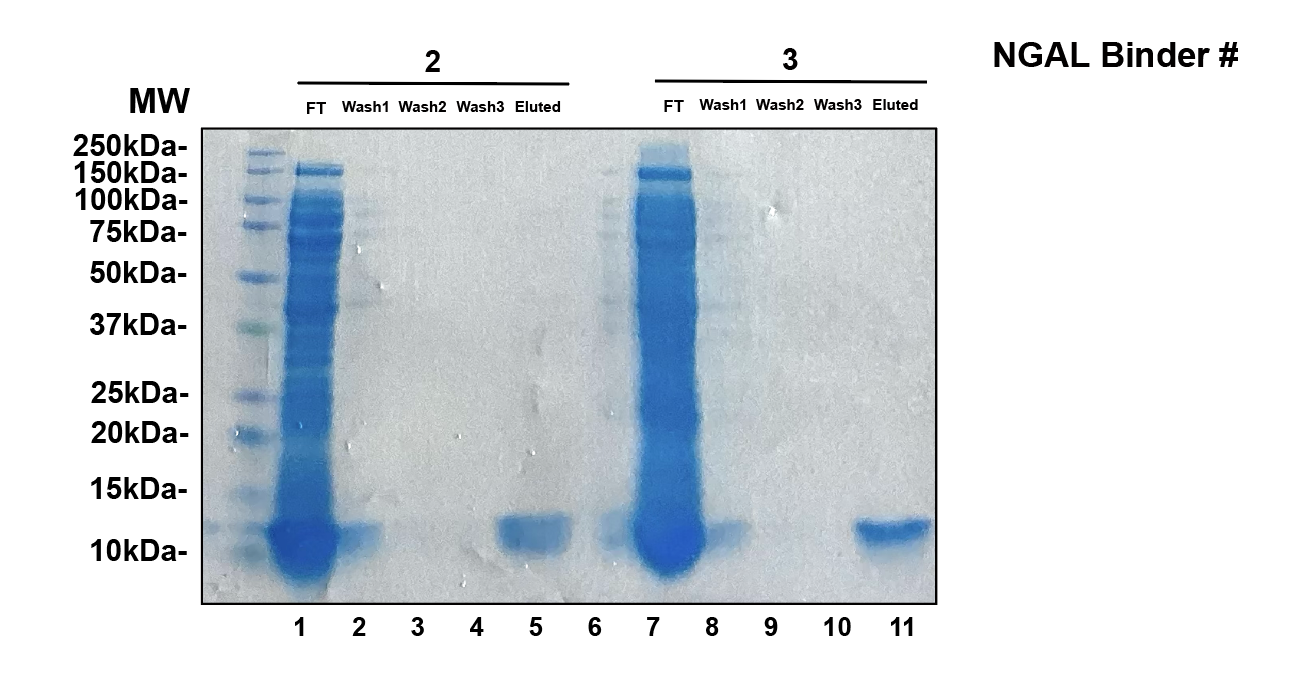

**Figure S2**. SDS-PAGE run to asses the miniprotein purity after production and purification.

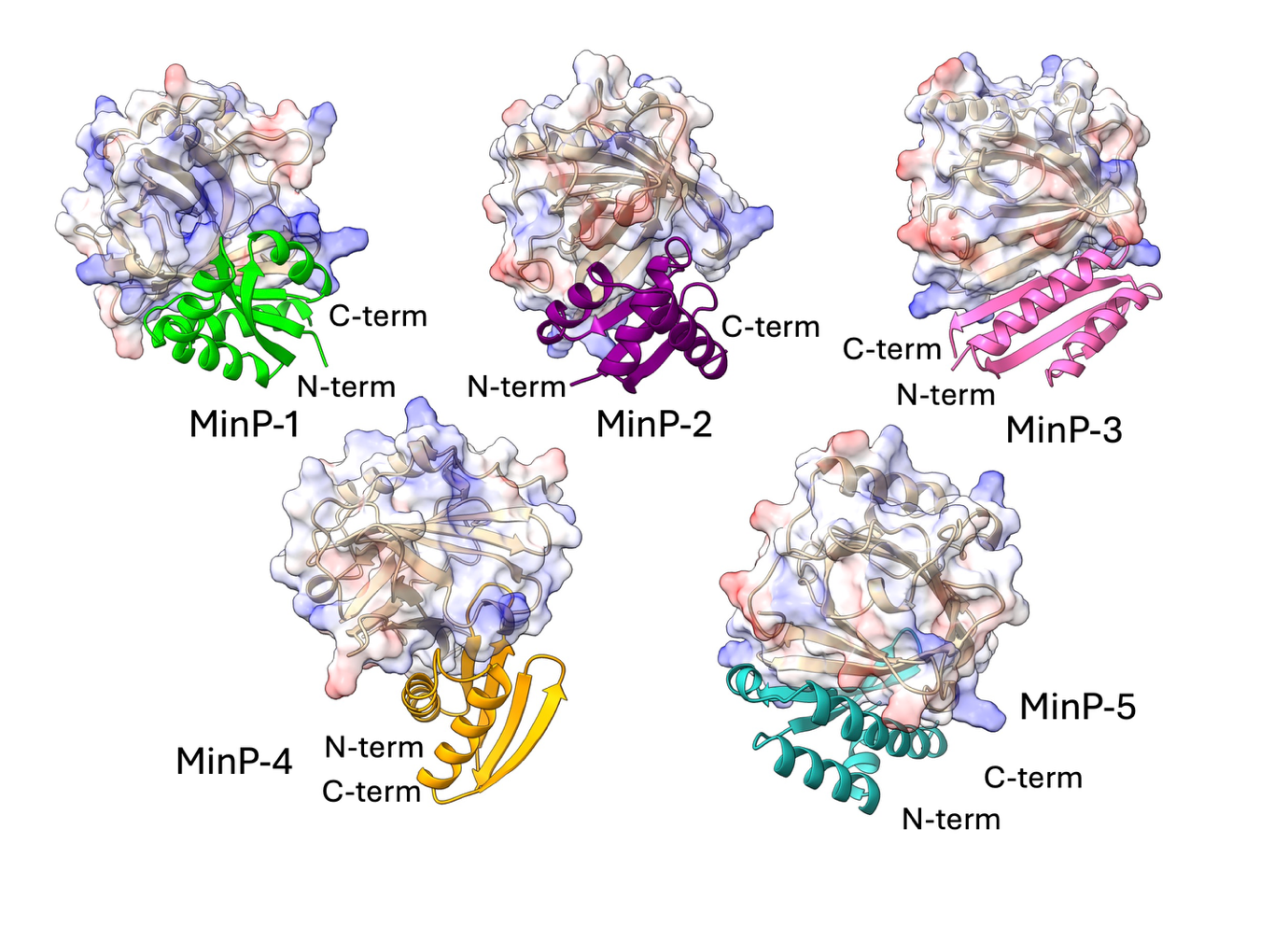
**Figure S3.** (B) Predicted structures of the complexes formed between LCN2 and the five selected miniproteins, as generated by AlphaFold2.

**Table S1**. Results alanine scanning on the residues located at the protein-protein binding interface for the MinP-2 in the complex with LCN2. The residue with a ΔΔG value higher than 0.5 kcal/mol are highlighted in yellow.

| **Residue Name** | **ΔΔG** |
| --- | --- |
| **LYS2** | 0.05 |
| **THR4** | 0.25 |
| **TYR6** | 0.19 |
| **VAL8** | 0.44 |
| **SER9** | 0.24 |
| **THR10** | 0.32 |
| **GLU15** | 0.1 |
| **LEU24** | 0.11 |
| **SER26** | 0.22 |
| **VAL29** | 0.77 |
| **TYR30** | 0.09 |
| **GLU31** | 0.3 |
| **TYR34** | 0.42 |
| **LEU36** | 0.08 |
| **ARG37** | 1.33 |
| **ILE38** | 0.67 |
| **SER39** | 0.35 |
| **VAL40** | 0.82 |
| **GLU43** | 0.09 |
| **ASP44** | 0.07 |
| **THR45** | 0.5 |
| **LYS46** | 0.05 |
| **ILE48** | 1.08 |
| **LEU49** | 0.63 |
| **ILE52** | 1.12 |
| **LYS53** | 0.3 |
| **ASN54** | 0.1 |

**Table S2.** Results alanine scanning on the residues located at the protein-protein binding interface for the MinP-2 in the complex with LCN2. The residue with a ΔΔG value higher than 0.5 kcal/mol are highlighted in yellow.

| **Residue Name** | **ΔΔG** |
| --- | --- |
| **ILE75** | 0.24 |
| **GLU77** | 0.48 |
| **LYS79** | 0.16 |
| **ASN85** | 0.45 |
| **THR87** | 0.21 |
| **VAL89** | 0.22 |
| **PHE91** | 0.23 |
| **ARG92** | 0.09 |
| **LYS94** | 0.05 |
| **LYS95** | 0.8 |
| **CYS96** | 0.36 |
| **ASP97** | 1.5 |
| **TYR98** | 3.16 |
| **TRP99** | 0.79 |
| **ILE100** | 1.91 |
| **ARG101** | 0.26 |
| **THR102** | 0.38 |
| **LEU114** | 0.08 |
| **ASN116** | 1.31 |
| **ILE117** | 0.13 |
| **SER119** | 0.23 |
| **TYR120** | 2.28 |
| **GLN194** | 0.06 |
| **CYS195** | 0.25 |
| **ILE196** | 0.16 |
| **ASP197** | 0.22 |
